# Overlaps in Brain Dynamic Functional Connectivity between Schizophrenia and Autism Spectrum Disorder

**DOI:** 10.1101/385146

**Authors:** Andry Andriamananjara, Rayan Muntari, Alessandro Crimi

**Affiliations:** AIMS-Rwanda, Kigali, Rwanda; AIMS-Ghana, CapeCoast, Ghana; University Hospital of Zurich, Zurich, Switzerland

**Keywords:** Brain imaging, Schizophrenia, Autism Spectrum Disorder, Connectome, Functional connectivity, FMRI

## Abstract

Schizophrenia and autism share some genotipic and phenotypic aspects as connectome miswiring and common cognitive deficits. Currently, there are no medical tests available for either disorders, and diagnostics for both of them include direct reports of relatives and clinical evaluation by a psychiatrist. Despite several medical imaging biomarkers have been proposed in the past, novel effective biomarkers or improvements of the existing ones is still need. This work proposes a dynamic functional connectome analysis combined with machine learning techniques to complement the present diagnostic procedure. We used the moving window technique to locate a set of dynamic functional connectivity states, and then use them as features to classify subjects as autism/schizophrenia or control. Moreover, by using dynamic functional connectivity measures we investigate the question whether those two disorders overlap, namely whether schizophrenia is part of the autism spectrum and which brain region could be involved in both disorders. The results reveal that both static and dynamic functional connectivity can be used to classify subjects with schizophrenia or autism. Lastly, some brain regions show similar functional flexibility in both autism and schizophrenia cohorts giving further possible proofs of their overlaps.

## 1. Introduction

Schizophrenia and autism spectrum disorder (ASD) are neurological disorders. Individuals who are affected by any of these disorders have difficulties with communication and with what is usually considered normal social behavior. Schizophrenia is a psychiatric disorder characterized by so-called positive symptoms as having delusion, hallucination, disordered thinking and speech and disorganized behavior [1]. Accompanying those, negative symptoms as reduced speaking, and reduced expression of emotions via facial expression or voice tone can occur [2]. ASD is a set of neuro-developmental disorders characterized by impaired social interaction and repetitive behaviors [3]. Among the expressed traits, deficits in nonverbal communicative behaviors used for social interaction can appear, as poor verbal and nonverbal communication, eye contact and body language deficits. In many cases subjects have also deficits in developing, maintaining, and understanding relationships, and social contexts [4]. Besides, the majority of ASD subjects fulfill diagnostic criteria for an anxiety disorder [5]. ASD severity is classified according to the level of support required by the ASD subject. [4].

Those disorders have been also studied by using magnetic resonance imaging (MRI) [6, 7]. Among the recent advances, connectomics is the most promising. A connectome is a comprehensive representation of the brain as a graph, where nodes are the brain regions, and edges represent connections either structural or functional among those brain regions [8]. The general spread miswiring in connectomes of subjects with either disorders has led the hypothesis that schizophrenia is part of the autism spectrum as the two disorders clearly overlap at some aspects [9]. Overall, schizophrenia is found to cause misconnections among brain regions [10] and ASD is identified with patterns of both high and low connectivity among brain regions [11, 3]. Both disorders are associated with decreases in inter-hemispheric connectivity. Particularly between frontal and posterior regions in the parietal lobe and occipital cortex [12, 13], and the corpus callous [14]. Recently, Yahata et al. identified a small number of connections which can discriminate ASD subjects from healthy control [15]. Mastrovito et al. compared atypical functional connectivity between ASD to typically developing children and schizophrenia to normal control, highlighting also common connectivity features between ASD and schizophrenia [16].

A new method is gaining attention in the study of brain activity: dynamic functional connectivity (dFC), an approach to analyze the functional connectivity from subsets of functional MRI (fMRI). This technique instead of using time series at once it uses parts of them defined by overlapping or non-overlapping windows of the overall series [17]. In this paper we focus on using dFC features with machine learning tools and novel graph metrics. More specifically we use support vector machine (SVM) for distinguishing either ASD or schizophrenia from control subjects. Moreover we quantify local dynamic differences and how they overlap between the two pathologies using the flexibility index of brain regions. Flexibility of each node/area corresponds to the number of times that it changes module allegiance (modularity) on time while we move across the dFC representation. A way to quantify changes in modularity, in such a dynamic network, is to perform clustering for each time-point independently and to count changes from one time point to another. However, computing cluster independently for individual time points has limitations as not taking into account the fact that same elements exist at different time points. Moreover, identifying the same labeling for each run can be cumbersome as many algorithms assign randomly labels at each run. Therefore, most common clustering approaches have not been available for time-dependent or multilayer networks, and ad hoc methods have been introduced to overcome those limitation [18, 19]. Mucha et al. developed a methodology, generalizing the determination of community structure via quality functions to multislice networks, which are defined by coupling multiple adjacency matrices as a generalized *Louvain modularity* [20]. For this reason this technique appears appropriate to cluster time dependent multilayer networks as dynamic FC matrices.

### 1.1. Functional Connectivity

*Functional Connectivity* (FC) is a measure of how temporally dependent processes interact. In the context of fMRI, these dependences is related to brain regions anatomically separated and their neuronal activation interact. In a functional connectome, edges are defined by the degree of association between regions during periods. Those associations can be determined by methods such as cross-correlations in the time or frequency domain, mutual information or spectral coherence [21]. *Static Functional Connectivity* is a global measure of functional connectivity of brain activity for the whole BOLD fMRI time series. *Dynamic Functional Connectivity* refers to the observed phenomenon that functional connectivity changes over time [22] even for resting-state acquisitions. It is sometimes referred as “time-varying” connectivity. Contrary to the previous notion that resting-state functional connectivity are stationary over time, studies revealed that when a brain is scanned over a period of time, it reveals a number of functional connectivity states which are susceptible to variations. Those have been also investigated with emphasis on matrix decompositions such as principal component analysis and independent component analysis [23]. Moreover, the nature of dynamic functional connectivity can be used to distinguish between healthy and afflicted brain [17, 24]. This technique has already been used in combination with SVM to classify subjects with traumatic brain injuries with high precision [25].

### 1.2. Moving Window Technique

Moving window analysis is a tool for analyzing the dynamic functional connectivity of brain activity detected from fMRI scan [26]. The concept of windowing is based on taking a time-window of fixed length and computing the functional connectivity of the data point inside of this time-window. After that, the window is moved with a fixed step, until the end of the time-series. Those steps can be defined to give overlapping or consecutive windows. The main parameters of window analysis are the window length and the window step. The results of any window analysis can largely influenced by the choice of these parameters [27]. However, there are no universally recognized values for these parameters and results can be just the consequence of over-fitting. The general rule is that the choice of window’s size should be large enough to permit robust estimation of functional connectivity and to resolve the lowest frequencies of interest in the signal, and yet small enough to detect potentially interesting transients [28, 27]. The choice of window length may depend on whether the time series changes rapidly. Most researchers use window length between 8 and 240s [24].

## 2. Method and Data

The present study seeks to use SVM to diagnose ASD/Schizophrenia and to look for overlaps between the two disorders. To achieve the first objective, for each subject the fMRI brain data was preprocessed and converted into FC matrix(matrices) by using windowing (fixed window for static or moving window for dynamic functional connectivity) and the Pearson correlation. The second objective is reached by using the flexibility score for different brain regions comparing either ASD or schizophrenia against control subjects, and then comparing the statistically significant results for between ASD and schizophrenia.

### 2.1. Data and Experimental Settings

We used two different datasets to perform the experiments. The ASD dataset is obtained from the publicly available ABIDE-II dataset [29]. More specifically the San Diego State University cohort comprising 54 subjects (31 ASD and 23 control), of age between 7 and 50 years, 72% male. For each subject, blood-oxygenation level dependent (BOLD) volumes were acquired in one 6:10 minute resting state scan consisting of 185 whole brain volumes (TR = 2, 000 milliseconds, TE = 30 milliseconds, flip angle = 90°, matrix size = 64 × 64 matrix, 3.4 × 3.4 × 3.4mm^3^ resolution, 42 axial slices covering the whole brain). The schizophrenia dataset is obtained from the the Center for Biomedical Research Excellence (COBRE) dataset [30]. The dataset included 146 subjects (2 discarded due to artifacts) ranging in age between 18 and 65 (70 with schizophrenia and 74 controls, 70% male). Practically, the two cohorts were matching per gender but only slightly per age as the COBRE cohort has a mean age of 36.5 years while the ABIDE cohort has a mean age of 22.65 years. BOLD volumes were obtained using TR = 2s, TE = 29ms, flip angle = 75°, 32 slices, voxel size = 3 × 3 × 4 mm^3^, matrix size = 64 × 64. Throughout the resting state scan, participants were instructed to relax, and to keep their eyes open and centered on a white fixation cross displayed on black background in the center of a screen, using a rear projection display.

### 2.2. Pre-processing and Connectome Construction

For both fMRI datasets data have been pre-processed according to a standard pipeline: motion correction, mean intensity subtraction, pass-band filtering with cutoff frequencies of [0.005-0.1 Hz] and skull removal. To account for potential noise from physiological processes such as cardiac and respiratory fluctuations, nine covariates of no interest have been identified for inclusion in our analyses [31]. To further reduce the effects of motion, compensation for frame-wise displacement has been carried out [32]. Linear registration has been applied between the Harvard-Oxford atlas [33] and the reference volume by using linear registration with 12 degrees of freedom. This atlas has been used during the experiments due to its small number of regions of interest (r=96) considering that the features used in the classification are the correlation matrices. In fact, the brain dataset produced a single 96 × 96 correlation matrix for each subject with rows and columns representing brain regions and the elements of the measures of association between the brain regions defined by the atlas. Due to the symmetry of correlation matrices, only the upper triangular parts were considered and flattened into 1D vectors for subsequent use as feature vector *x_i_*. For any given symmetric matrix of dimension *N* × *N*, the upper triangular gives a total of *N*(*N* − 1)/2 elements, which for the static FC means a feature vector of 4560 elements. The length of the windows has been chosen in nested cross-validation manner identifying the the shape leading the highest classification but simpler configuration given by less windows used. Namely 30 time points and non-overlapping windows, generating 5 time-windows. This values have been used for both dataset for consistency. As all the time series of the schizophrenia dataset were 150 time points, and the time series for the ASD dataset were 180 time points (the last 30 times points were discarded for consistency).

### 2.3. Case-Control Classification

The used features for the SVM-based classification are the dFC matrices. In our experiments, functional connectivity is defined by using the Pearson correlation between variables *A* and *B* being brain region as

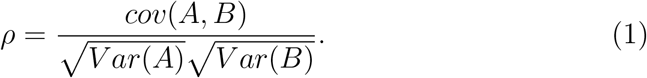

In windowing analysis, we define correlation time series 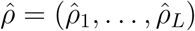 over a functional brain data by successively moving, at a constant rate, a window of predetermined length and shape over the data. We compared results obtained using one static FC matrix per subject with results using several dFC matrices per subject. Dynamic functional analysis was performed using using the concatenation of the 1D vectors. The dataset were labeled accordingly and used to train and test the selected SVM model. The validity of the model was evaluated using leave-one-out cross validation and Receiver Operating Characteristic (ROC) curve.

Here we review the main concept of SVM [34]. SVM is a supervised learning method which constructs a hyperplane or set of hyperplanes in a high-or infinite-dimensional space used for classification. Considering the problem for binary classification, such as ASD versus control, we have two classes of subjects (samples) and we want to separate (classify) them. The data are given as *x*_1_, *x*_2_, …, *x_m_* ∈ *X, y*_1_, *y*_2_, …, *y_m_* {±1}. Such that *X* is a non-empty set in the domain of *x_i_* features, and *y_i_* are the targets or labels. We assume that the data are into the Hilbert product space 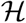 to allow us to measure the distance of the vectors in that space. Among all hyperplanes we seek for the optimal separable hyperplane that maximize the distance between the data points (vectors). This can be achieved by using the hyperplane decision function

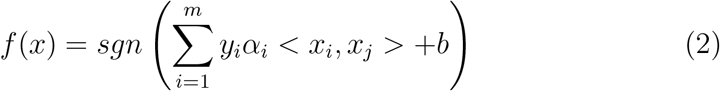

where *α_i_* are Lagranger multiplier and *b* is a threshold which can be computed by convex optimization [34].

### 2.4. Functional Connectivity Differences

As we are mostly interested in dynamic functional connectivity, we used a metric quantifying changes in it. The flexibility index for each node of a connectome defined at different time points is defined as the number of times that it changes community allegiance, normalized by the total possible number of changes. We consider changes in functional connectivity possible only between consecutive time points. The flexibility index has been introduced by Bassett et al. [35] to quantify changes in connectomes related to learning at different time points. Biologically, network flexibility could be driven by physiological processes that facilitate the participation of cortical regions in multiple functional communities (clusters) [35]. As communities organization change smoothly with time, the flexibility index displays coherent temporal dependence also in dynamic functional changes within the same session. Fig. 1 gives a schematic representation of a multilayer graph with two nodes that change community allegiance twice along time. Our rationale is that ASD and schizophrenia subjects have some brain regions with different flexibility from healthy control subjects due to their connectome miswiring. Moreover, we are interested in seeing if some of these differences are shared between ASD and schizophrenia subjects. To validate this, we assessed initially the number of clusters in all our dFC connectomes using the eigengap techniques related to spectral clustering [36] as a mean eigengap for all connectomes. The eigengap of a linear operator is the the difference between two successive eigenvalues, where eigenvalues are sorted in ascending order. In the context of spectral clustering, we define a Laplacian matrix **L** = **D** – **A** with **D** the diagonal matrix containing the degrees of each node in the graph/connectome, and **A** the adjacency matrix of the graph in our case defined by the Pearson correlation. Then, we perform an eigendecomposition, and we denote the ordered eigenvalues of **L** matrix as *λ*_0_ ≥ *λ*_1_ ≥ · · · ≥ *λ_r_*. The eigengap is largest difference in absolute value of two consecutive eigenvalues as *λ_i_* − *λ*_*i*+1_. Therefore, we computed the number of clusters as a mean *μ* across all the connectomes for those largest value

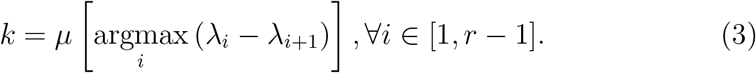

**Figure 1:**
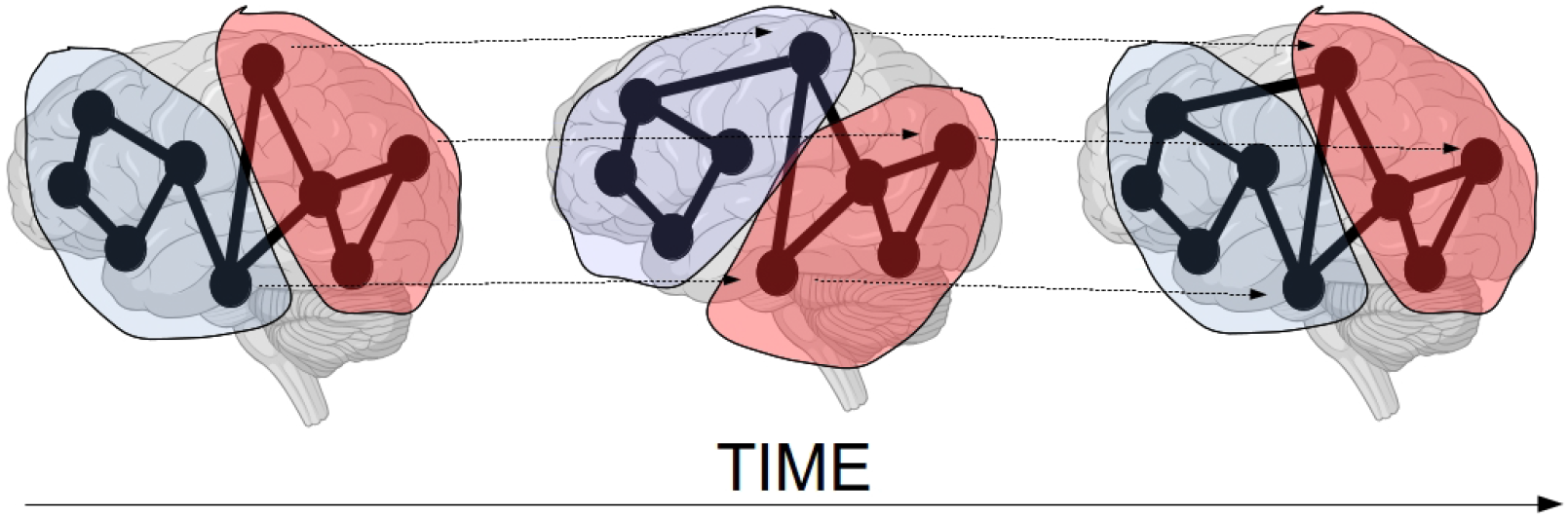
Schematic representation of multilayer graph with two nodes which change community across time twice. The dashed lines highlight the corresponding brain regions, which are also intracommunity edges. Each time point is a slice/layer of the multilayer network.

Nevertheless, to avoid clustering techniques which do not take into account the multi-layer nature of dynamic FC matrices, we resorted to the generalized Louvain modularity which also include the cluster number discovery in its implementation [20]. More specifically, for an undirected weighted or binary graph the multislice modularity can be defined as

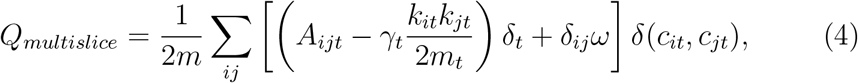

where *A_ijt_* is the weight of the edge connecting between nodes *i* and *j* at time t from the adjacency matrix **A**, *k_it_* and *k_jt_* are the sums of weights of the edges connected to node *i* and *j* at *t* respectively, 2*m_t_* and 2*m* are the sum of all of the edge weights in the graph at time/slice *t* and for all time/slice respectively, *c_it_* and *c_jt_* are the communities of nodes *i* and *j* at *t, δ*s are delta functions, *ω* is the interslice coupling which represents the direct count of the intracommunity edge weights, and *γ_t_* is the spatial resolution [37]. In our experiments we set *ω* = 0.1 and *γ_t_* = 1 following the suggested used parameters by Mucha et al. [20] without optimizing them.

Once clustering has been carried out and module allegiance for each brain region across time is known. Changes across time can be defined by the *flexibility* of a node *f_i_* normalized by the total number of changes that were possible. Given all *f_i_* index for each brain region and subjects, a statistical framework based on two-tail t-test was used to find statistical difference between groups (ASD vs control, schizophrenia vs control, and ASD vs schizophrenia). T-test values were then converted into p-values to check which are statistically significant under the threshold *α* = 0.005. Afterwards the Benjamini & Hochberg procedure for controlling the false discovery rate [38] has been conducted and the adjusted p-values are reported in Table 1 and Table 2.

**Table 1:** ROIs found statistically significant comparing schizophrenia to healthy control subjects and their p-values. The ROIs significant also using the ASD datasets are highlighted in italic.

| # | ROI | p-value Schizophrenia-Control |
| --- | --- | --- |
| 1 | Superior Frontal Gyrus Left | $< 0.001$ |
| 2 | Superior Temporal Gyrus, anterior division Left | $< 0.001$ |
| 3 | *Superior Temporal Gyrus, posterior division Left* | *0.0013* |
| 4 | *Middle Temporal Gyrus, posterior division Right* | $< 0.001$ |
| 5 | Frontal Medial Cortex Left | $< 0.001$ |
| 6 | Cingulate Cortex anterior Right | 0.0013 |
| 7 | *Cingulate Cortex posterior Left* | *0.0010* |

**Table 2:**
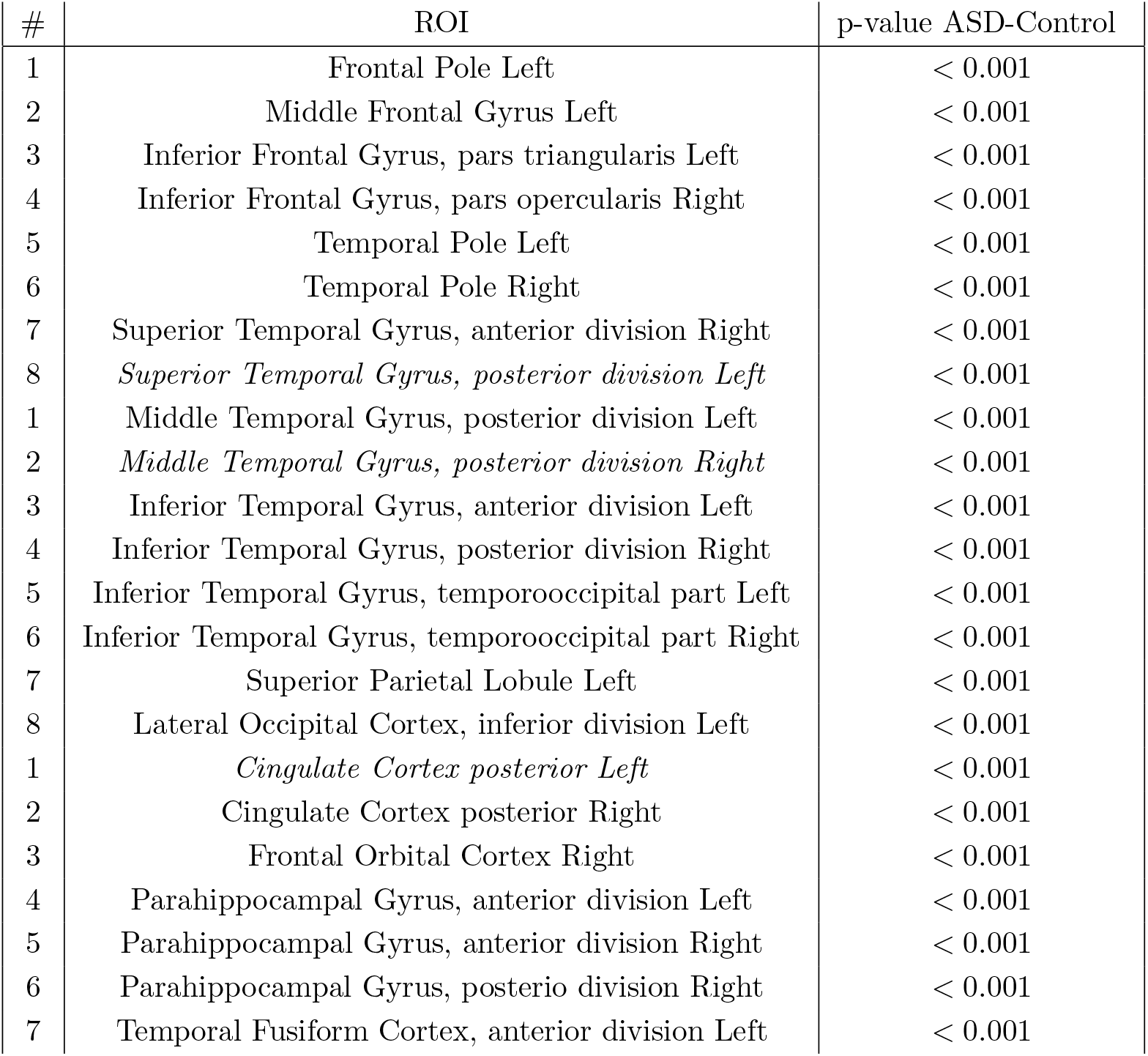
ROIs found statistically significant comparing ASD to healthy control subjects and their p-values. The ROIs significant also using the schizophrenia datasets are highlighted in italic.

| # | ROI | p-value ASD-Control |
| --- | --- | --- |
| 1 | Frontal Pole Left | < 0.001 |
| 2 | Middle Frontal Gyrus Left | < 0.001 |
| 3 | Inferior Frontal Gyrus, pars triangularis Left | < 0.001 |
| 4 | Inferior Frontal Gyrus, pars opercularis Right | < 0.001 |
| 5 | Temporal Pole Left | < 0.001 |
| 6 | Temporal Pole Right | < 0.001 |
| 7 | Superior Temporal Gyrus, anterior division Right | < 0.001 |
| 8 | *Superior Temporal Gyrus, posterior division Left* | < 0.001 |
| 1 | Middle Temporal Gyrus, posterior division Left | < 0.001 |
| 2 | *Middle Temporal Gyrus, posterior division Right* | < 0.001 |
| 3 | Inferior Temporal Gyrus, anterior division Left | < 0.001 |
| 4 | Inferior Temporal Gyrus, posterior division Right | < 0.001 |
| 5 | Inferior Temporal Gyrus, temporooccipital part Left | < 0.001 |
| 6 | Inferior Temporal Gyrus, temporooccipital part Right | < 0.001 |
| 7 | Superior Parietal Lobule Left | < 0.001 |
| 8 | Lateral Occipital Cortex, inferior division Left | < 0.001 |
| 1 | *Cingulate Cortex posterior Left* | < 0.001 |
| 2 | Cingulate Cortex posterior Right | < 0.001 |
| 3 | Frontal Orbital Cortex Right | < 0.001 |
| 4 | Parahippocampal Gyrus, anterior division Left | < 0.001 |
| 5 | Parahippocampal Gyrus, anterior division Right | < 0.001 |
| 6 | Parahippocampal Gyrus, posterior division Right | < 0.001 |
| 7 | Temporal Fusiform Cortex, anterior division Left | < 0.001 |

Moreover, we looked at static and dynamic FC differences using the network based statistics (NBS) [10]. NBS is a nonparametric statistical test used to identify connections within connectivity matrices which are statistically significant different between two distinct populations [10]. In practice, the NBS checks the family-wise error rate, where the null hypothesis is tested independently at each of the edges. This is achieved performing a two-sample t-test at each edge independently using the values of connectivity. The tests are repeated *h* times, each time randomly permuting members of the two populations. In our experiments we set *h* = 1000.

As noticed empirically that the NBS was producing about 300 connection using a p-value threshold related to the t-test of *α* = 0.05 (proportionally in agreement with Mastrovito et al. [16]), we lowered the NBS threshold to *α* = 0.01 to allow visual inspection of those results.

Most of the used code and the pre-processed time series are available online at the URL *https://github.com/alecrimi/dyfunconnclustering*.

## 3. Results

We carried out the experiments in a nested leave-one-out fashion. It was noted that varying the shape of the non-overlapping windows has an impact on the classification performed. Generally, dynamic FC matrices led to better classification than static FC. Figure 2 depicts all AUC values for both datasets for both static FC (1 window), and different windows size (2 or more windows) for the dynamic FC. We consider 13 windows as the lower limit as this setting requires windows of 10 time-points which is expected to be the minimum considering the cut-off frequency defined in the preprocessing of 0.1 Hz [27]. It can be noted that several windowing condition can lead to similar results.

**Figure 2:**
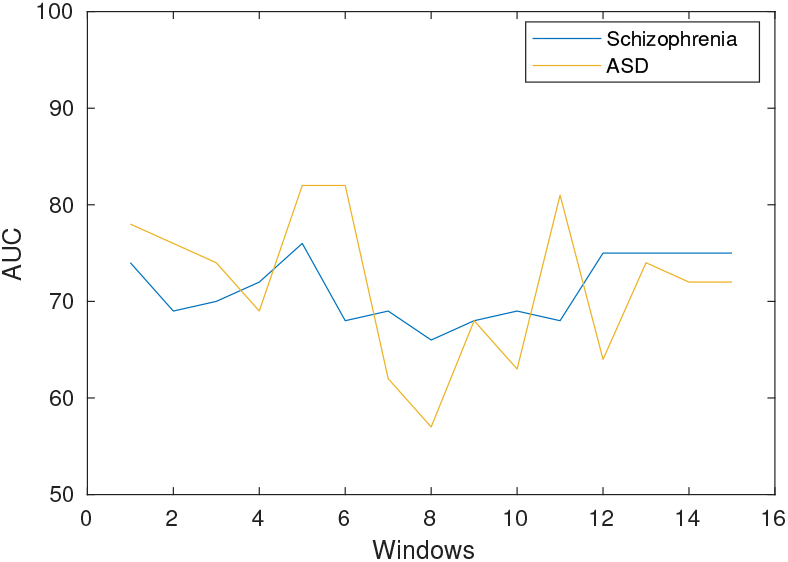
AUC varying according to the number of windows used and therefore the shape of the windows. The static FC is represented by the value window=1.

**Figure 3:**
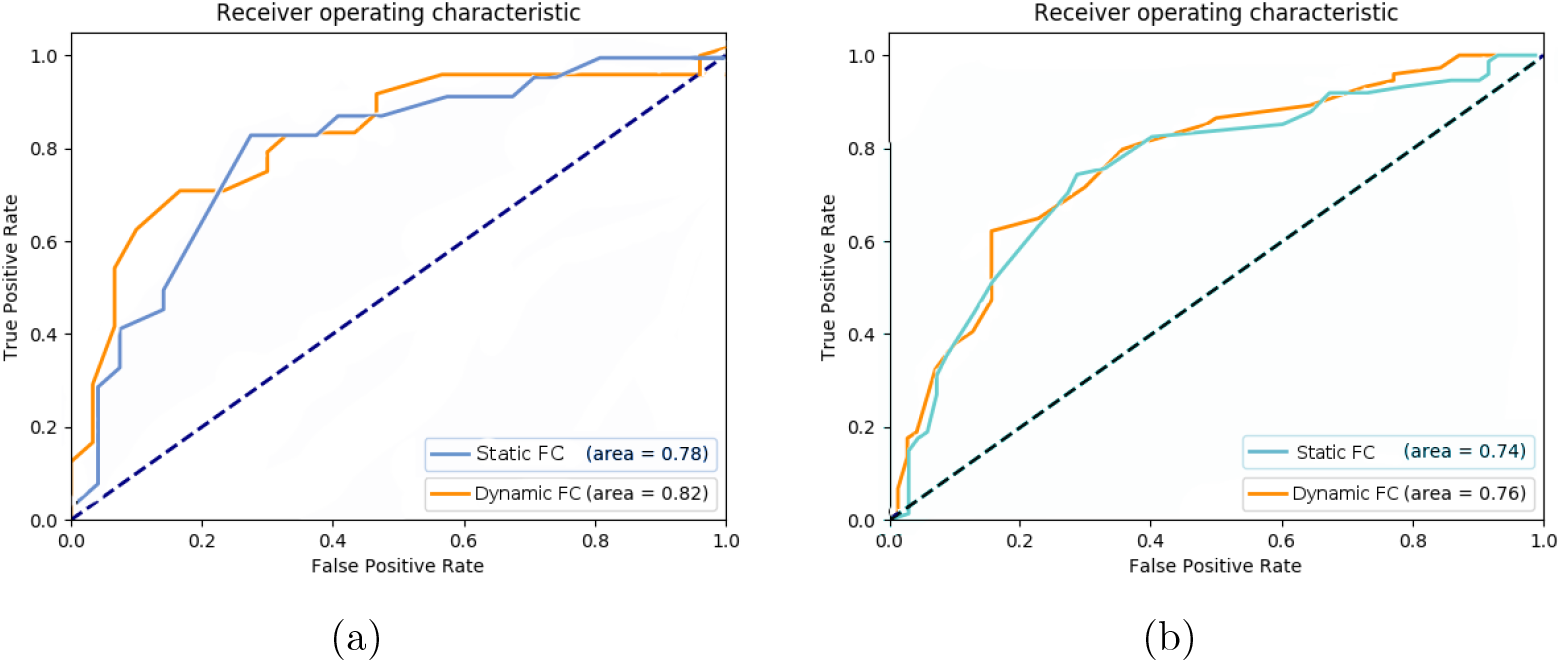
ROCs for static functional connectivity analysis. (a) ASD and schizophrenia (b)

Ultimately, for the ASD dataset the resulting area under the ROC (AU-ROC) was 0.78 using the static FC matrix, while the AUROC value was 0.82 using the dynamic FC matrices as shown in Figure 5 (a).

For the schizophrenia dataset the AUC was 0.74 if the static FC features were used, and 0.76 if the dynamic FC features with window size 30 were used, as shown in Figure 5 (b).

Surprisingly both the eigengap and the optimization of the Louvain modularity identified the same number of clusters as *k* = 4. The statistically significant areas (*α* = 0.005) according to the flexibility index are reported for schizophrenia and ASD dataset respectively in Table 1 and Table 2. The experiments with both datasets highlighted the posterior cingulate cortex, and parts of the superior and middle temporal gyrus as different regions from the flexibility point of view.

The static local differences detected by using NBS however were not directly comparable between the ASD and schizophrenia cohorts, apart the presence of DMN related connections to the posterior cingulate gyrus (CGp) and medial frontal cortex (FMC) as depicted in Figure 4. The dynamic local differences detected by using NBS were different from the static ones. Nevertheless, CGp and superior temporal gyrus left (T1p.L), paracingulate gyrus right (PAC.R) and temporal occipital fusiform cortex (TOF.L) were the main nodes with statistically different connections as shown in Figure 5.

**Figure 4:**
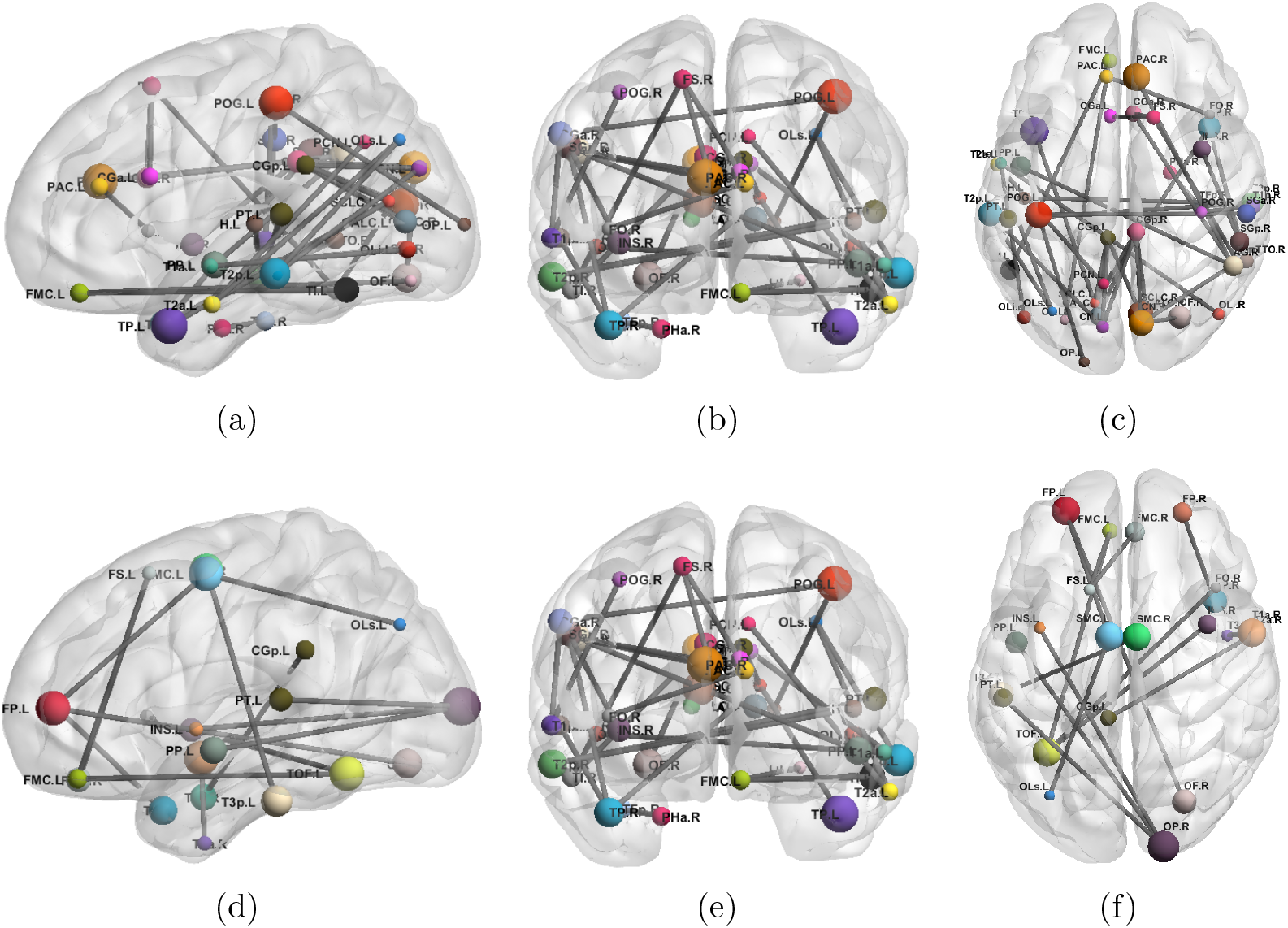
Statistically significant connection detected on the static FC matrices respectively for the schizophrenia (a) sagittal, (b) coronal, and (c) axial view. Statistically significant connection of the ASD cohort against respective control (d) sagittal, (e) coronal, and (f) axial view. The grey lines represents the discrininamt connections for the static FC matrices.

**Figure 5:**
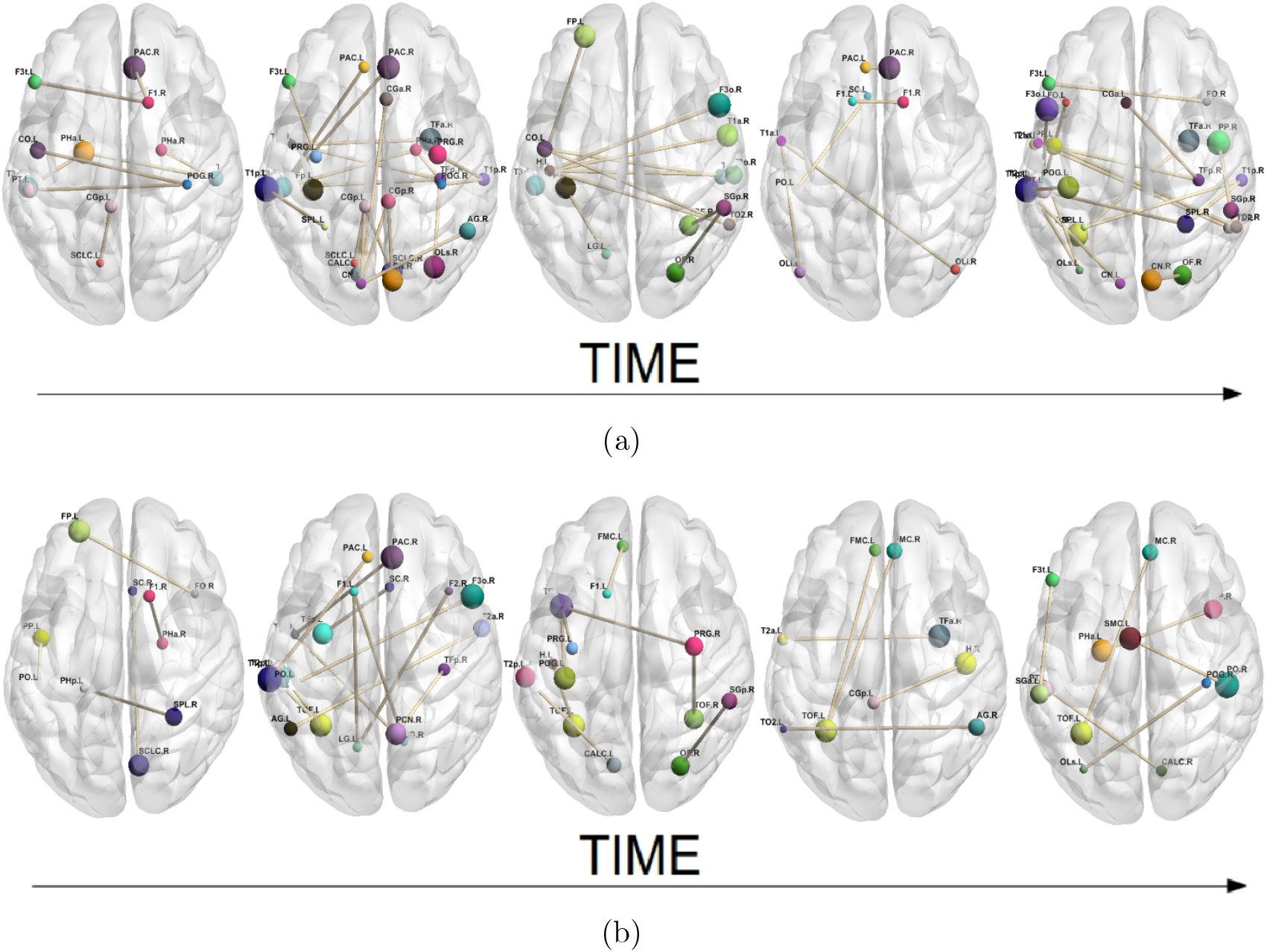
Discriminant connections detected by NBS for different windows ordered temporally from left to right. (a) Schizophreana and (b) ASD. The brown lines represent the discriminant connections for the dynamic FC matrices.

## 4. Discussions

Several studies support the idea of continuum between schizophrenia and ASD. Previous investigations showed that subjects with high-functioning ASD shared similar but more severe impairments in verbal theory of mind than schizophrenia patients [39]. More recently the analysis of the co-atrophy network of schizophrenia, ASD, and obsessive compulsive disorder showed that alterations in certain grey matter ROIs appear to be statistically related to alterations of other grey matter ROIs consistently in all three disorders [40]. From a diagnostic point of view, it could be useful to identify biomarkers for psychiatric disorders such as ASD and schizophrenia. It has been already hypothesized that schizophrenia subjects present higher flexibility indices than control in their connectomes probably due to mechanism modulated by N-methyl-D-aspartate receptors [41]. In line with this work, our experiments with both datasets highlighted the cingulate gyrus as a different region from the flexibility point of view. In other words, the cingulate gyrus and middle temporal gyrus for both ASD and schizophrenia subjects manifest an altered reconfiguration compared to control subjects. This can be explained by the fact that those regions are among the main hubs of the default mode network (DMN) which is known to be different for both ASD and schizophrenia subjects compared to healthy controls [16]. Many studies have identified the DMN as a collection of areas that are structural and functional hubs related to supplementary motor-areas, frontal eye fields involved in control of visual attention [42, 43]. Although the two disorders are known to exhibit significant changes in connectivity in the DMN, the two disorders display different connectivity alterations. ASD subjects exhibit a greater proportion of within-network changes in the DMN and reduced connectivity between DMN and language areas. Conversely, schizophrenia changes between DMN and language areas are increased in connectivity [44].

Despite the local differences identified by NBS were not directly comparable between the ASD and schizophrenia cohorts, the detected discriminant connections individually for ASD and schizophrenia are in line with connections detected by other works [15, 45, 16]. Nevertheless, the general spread miswiring for both disorders yet suggests the hypothesis that schizophrenia is part of the autism spectrum as the two disorders clearly overlap at some aspects [9]. Further hypothesis also relate attention deficit hyperactivity disorder (ADHD) to the spectrum [46, 47]. Additional investigations relating ADHD to ASD or schizophrenia could be interesting but they are beyond the purpose of this paper.

Alternatively to Pearson correlation, partial correlation measure the strength of the relationship between two variables, while after ruling out third-party effects [48]. Despite partial correlation would be expected to be more reliable, Smith et al. showed that both correlation methods provide reliable estimates of functional connections, but Pearson correlation outperforms partial correlation when the number of nodes in brain networks significantly increased [49]. Moreover, in a comprehensive comparison Pearson correlation-based brain networks had the most reliable topological properties compared to those estimated by using partial correlation and an atlas with similar parcellation to those used in our experiments [50].

A limitation of the study is given by the two cohorts (ASD/control and schizophrenia/control) being acquired by different centers with slightly different protocols. Despite it remains unclear whether potential confounding factors are introduced by using different scanners. Some studies showed that data can be pooled from different scanners without corroding the results as for certain measurement of Alzheimer [51], and in fact both datasets have been used by Mastrovito et al. also pooling them [16]. However, we cannot completely rule out confounding factors.

In our experiments the classification using the dynamic FC features outperformed the classification using the static FC features, though this difference might not necessarily be statistically significant. Comparing the results to state-of-art method, Yahatama et al. used static FC features and a sparse logistic regression classifier to discriminate ASD from control subjects obtaining an AUC = 0.93 on a Japanese cohort and AUC = 0.74 on the ABIDE dataset, but lower results on an another schizophrenia dataset [15]. Therefore, it seems that performances are strictly related to the cohort in use. Furthermore, Mastrovito et al. using static FC features on similar datasets to those used in the proposed approach, obtained accuracy ranging from 33% to 83% on the ASD dataset, and from 35% to 80% for the schizophrenia dataset varying the used features [16]. Therefore, this suggests that a further improvement could be obtained using dimensionality reduction or independent component analysis, as shown in [52]. The high dimensionality of the features has also influenced the choice of the Harvard-Oxford atlas. Indeed, atlases more specific for functional data exist [53]. However, those are generally with higher number of ROIs compared to the Harvard-Oxford leading to even higher dimensional features. We have avoided the use of those highly parcellated atlas as it would have complicated even more the framework but it is nevertheless acknowledged as a limitation.

## 5. Conclusions

This work confirm the potentiality of using machine learning techniques - as SVM - jointly to dynamic functional connectivity features as a further tool for discriminating both ASD and Schizophrenia from healthy subjects. Both static and dynamic functional connectivity can be used as features for this discrimination. Nevertheless, rather than proposing optimal classification for those two disorders, the focus was on their similarities, where the fact that classification by machine learning and functional connectivity is possible is only one aspect. Indeed, flexibility index and NBS highlighted the impact of the posterior cingulate gyrus and superior temporal gyrus left for both disorders compared to healthy control. This can be a starting point to further investigate similarities and overlaps between those and other disorders.

### Statements

No actual or potential conflict of interest including any financial, personal or other relationships with other people or organizations within three years of beginning the submitted work exists.

